## Supplementary Figures for "State-specific morphological deformations of the lipid bilayer explain mechanosensitive gating of MscS ion channels"

Yein Christina Park[1], Bharat Reddy[2], Navid Bavi[2],
Eduardo Perozo[2]* and José D. Faraldo-Gómez[1]*

[1]Theoretical Molecular Biophysics Laboratory
National Heart, Lung and Blood Institute
National Institutes of Health, Bethesda, MD

[2]Department of Biochemistry and Molecular Biology
University of Chicago, Chicago, IL

*Correspondence should be addressed to:

January 16th, 2023

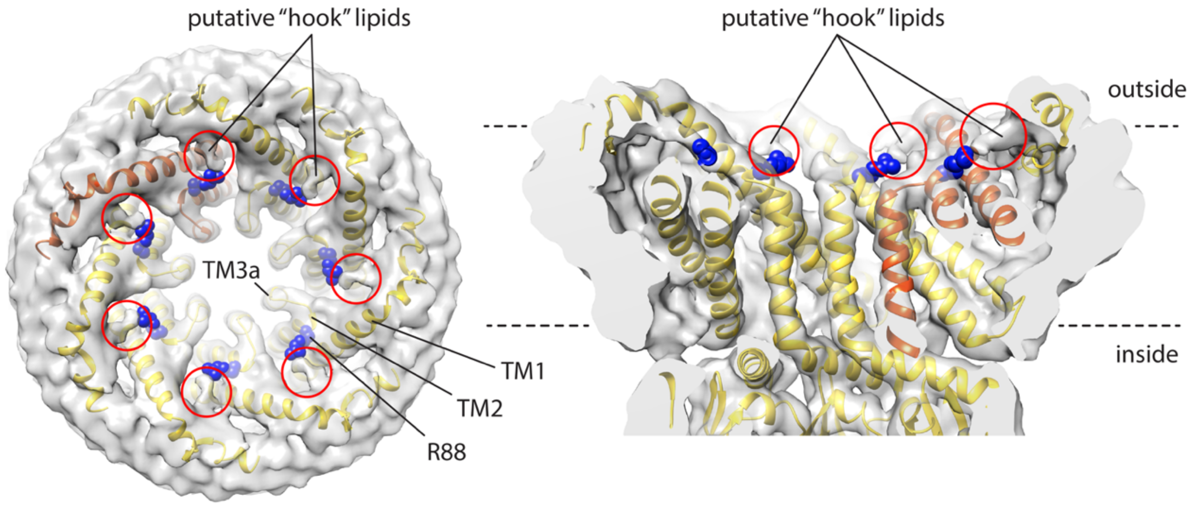


**Figure 2 – Figure Supplement 1.** Cryo-EM map and structural model of open-state MscS in PC14:1 lipid nanodiscs, highlighting putative sites of lipid interaction atop the C-terminus of TM2, involving residue R88. Lipids at these sites were more clearly discerned in the closed state in PC18:1 nanodiscs [17], and referred to as 'hook' lipids.

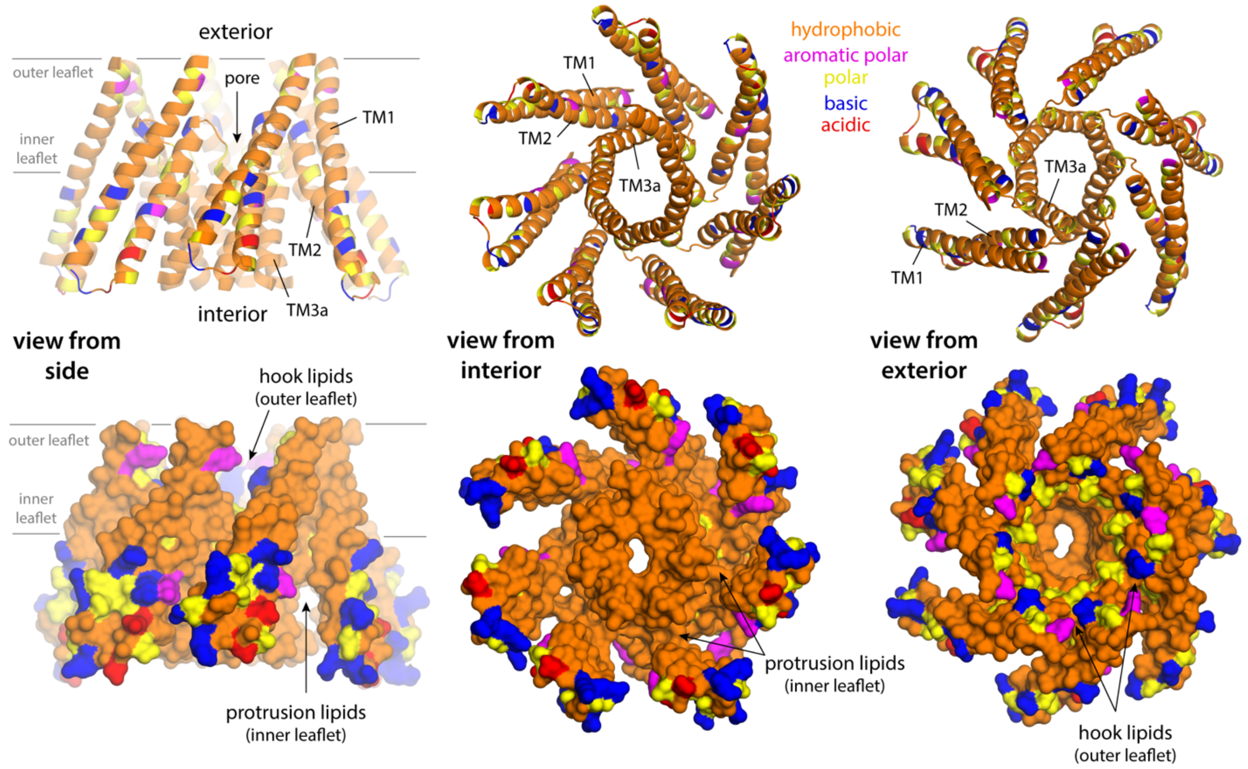


**Figure 4 – Figure Supplement 1. Lipid solvation of hydrophobic cavities outside the membrane drives the formation of inner-leaflet protrusions in closed-state MscS.** The figure shows three different views of a fragment of the closed MscS structure comprising TM1, TM2 and TM3a (from left to right), in two alternative representations (top and bottom). Residues are colored according to type, as indicated. The location of the inner leaflet protrusions under the TM1-TM2 hairpin is indicated; the site where the so-called "hook" lipids are observed is also indicated. The conformation of the channel that is represented is that in the all-atom snapshot shown in **Figure 5A**.

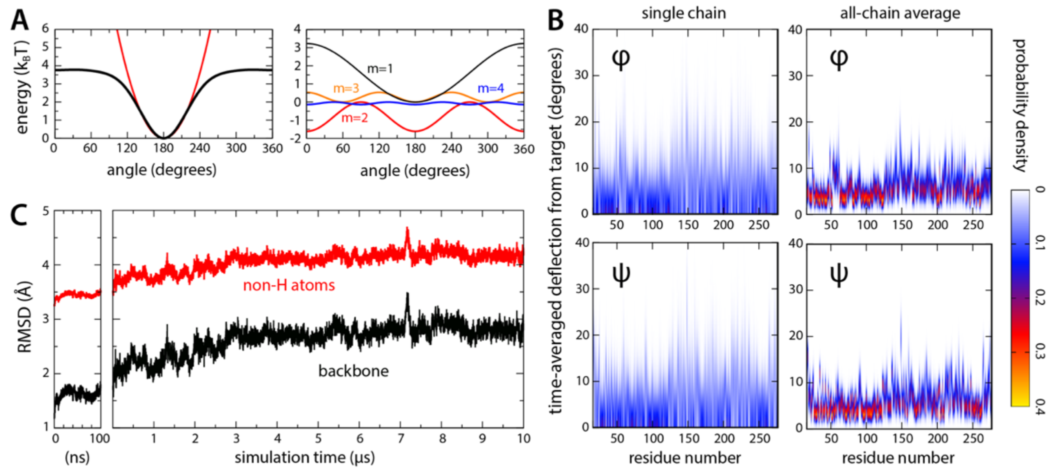


**Figure 4 – Figure Supplement 2. Relaxation of closed-state MscS structure in all-atom simulations.** (**A**) To preclude large-scale changes in fold that might develop in the 10-μs timescale due to cumulative forcefield inaccuracies, a restraining potential was applied to all $\phi$ and $\psi$ angles in the channel backbone, of the form $U(\theta_t) = k \sum_{m=1}^{m=6} (-1)^m \left[1 + \cos\left(m\theta_t - m\left(\theta_{expt} - 180\right)\right)\right] / m!$ , where $\theta_t$ is the value of either dihedral at time $t$ in the simulation, $\theta_{expt}$ denotes the value in the experimental structure, and $k$ equals 4 kJ/mol (see Materials and Methods). The black curve in the left plot in (A) exemplifies this potential for $\theta_{obs}$ = 180; for comparison, a harmonic potential $U'(\theta_t) = k\ (\theta_t - \ \theta_{obs})^2$ is superimposed (red line), where $k = 0.0025$ kJ/mol/deg$^2$. The right plot in (A) shows each of the first four terms contributing to $U(\theta_t)$. (**B**) For each residue in the channel, observed deflections in $\phi$ and $\psi$ during the simulation relative to the corresponding values in the experimental structure, individually quantified as probability density distributions. The left plots show data for a single chain in the heptamer, selected at random; the right plots show a global analysis for all seven chains. (**C**) RMS difference between simulated and experimental channel structure, as a function of simulation time. Data is provided for the backbone atoms only, and for backbone and sidechains, excluding hydrogen atoms. The RMS difference calculation was preceded by least-squares superposition of the protein backbone in the snapshot considered and in the experimental structure. The plot on the left shows data for the initial equilibration of the simulation system, carried out using NAMD; the plot on the right shows data for the ANTON2 simulation (see Materials and Methods).

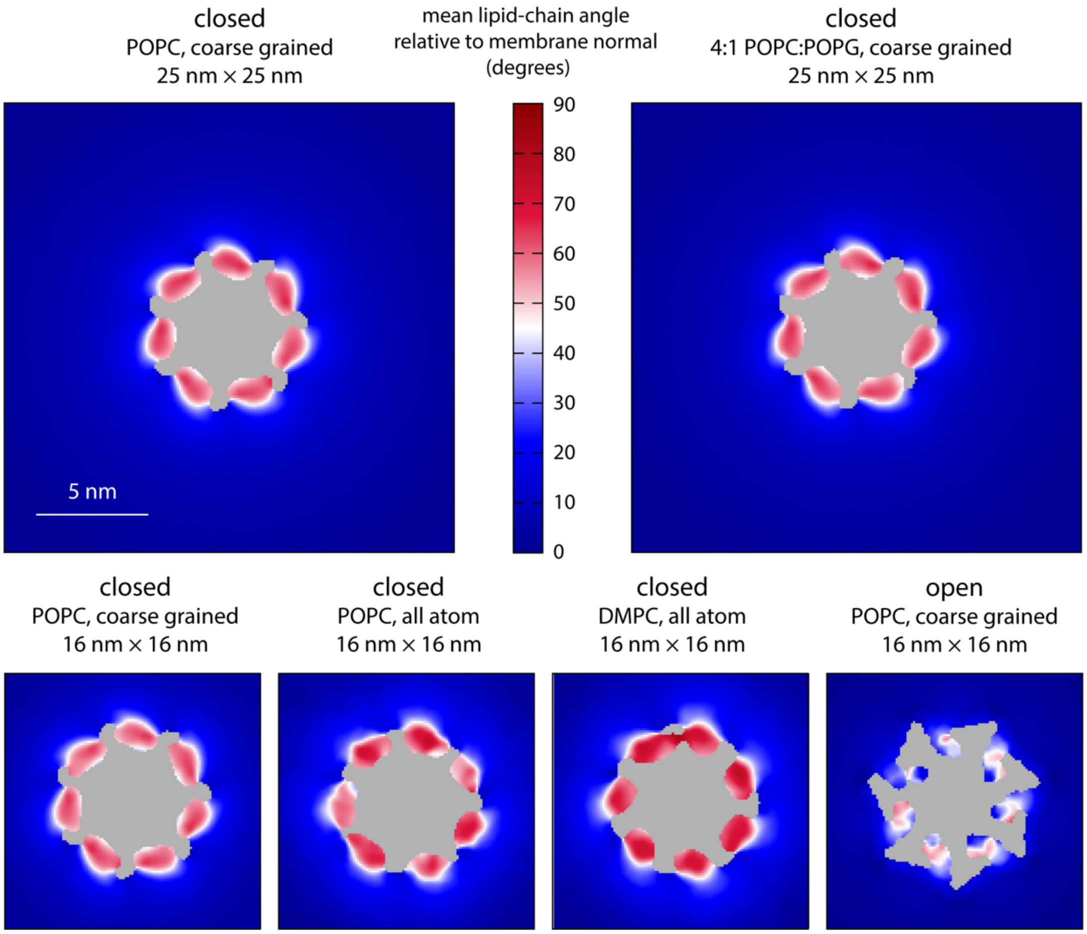


**Figure 5 – Figure Supplement 1. Molecular structure of the membrane perturbations induced by MscS in the closed state.** The figure reports results for multiple simulations of closed- and open-state MscS in membranes of different size and lipid composition using either coarse-grained or all-atom representations (**Table 1**). Specifically, the plots map the characteristic tilt of the lipid alkyl chains across the inner leaflet of the bilayer, relative to the membrane perpendicular, derived from analysis and averaging of instantaneous lipid configurations. The central areas (gray) are occupied by protein or solvent filling the central pore. The maps are derived from 20 μs of trajectory data for each of the coarse-grained systems and at least 8 μs of trajectory data for the all-atom systems.

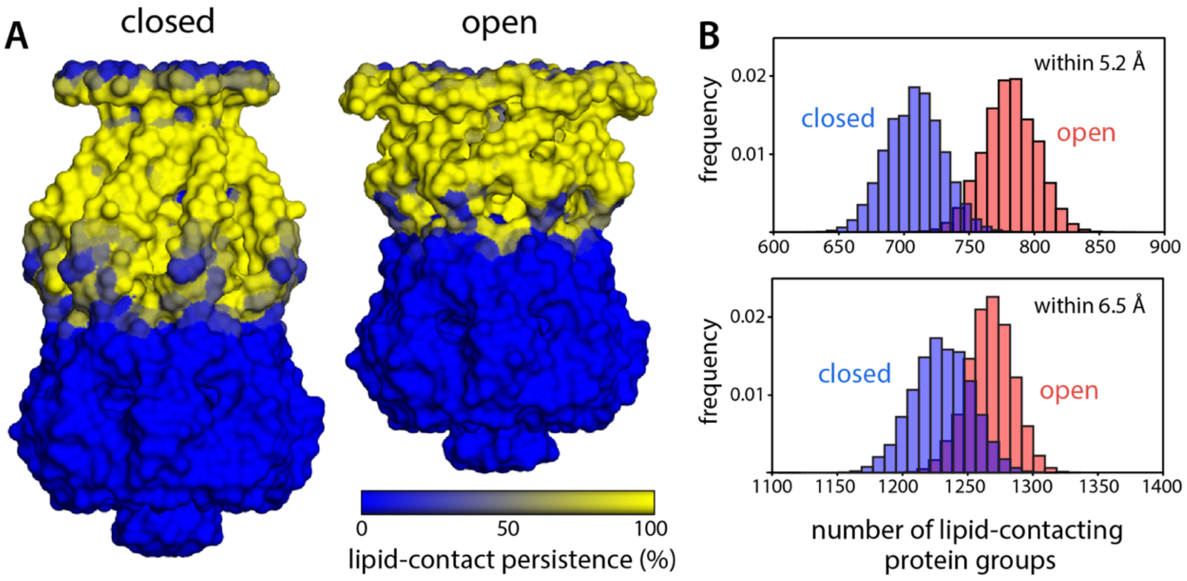


**Figure 8 – Figure Supplement 1. Protein-lipid interfacial area is largely unchanged during gating.** (**A**) To evaluate the area of the protein surface exposed to the membrane, we quantified the number of coarse-grained (CG) particles in the channel within 6.5 Å of any lipid particle, for each snapshot of the MD trajectories calculated for closed and open MscS in a POPC lipid bilayer. Note that each CG particle approximately represents one chemical group in protein sidechains and backbone, and that a distance of 6.5 Å implies either a direct or close-range contact, as the typical radius of the CG particles is 2.6 Å. The results from this analysis are mapped on the channel surface, for either functional state, coloring each CG particle according to the persistence of their exposure to the lipid bilayer over time. (**B**) Histograms quantifying the variability in number of CG particles in the channel that are exposed to the lipid bilayer across different snapshots within the same trajectory. For completeness, data are shown for a distance threshold of 6.5 Å, as in panel (A), as well as for 5.2 Å, which implies direct contact. For threshold 5.2 Å, the number of lipid-contacting protein groups is 708.2 ± 21.4, while in the open state this number is 781.3 ± 20.1, reflecting a 10.3% increase. For threshold 6.5 Å, the corresponding values are 1230.3 ± 22.5 and 1267.9 ± 17.6, reflecting a 3.0% difference.

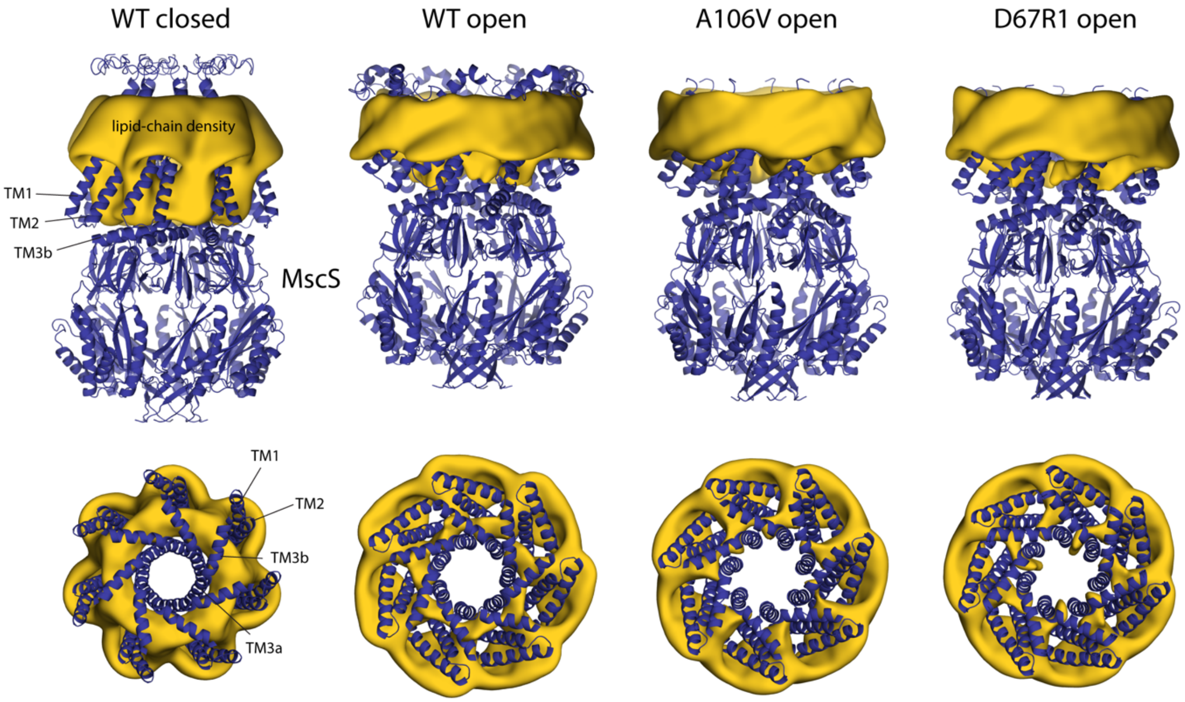


**Figure 8 – Figure Supplement 2. Changes in membrane morphology upon gating of MscS.** The figure summarizes results from simulations of alternative open conformations of wild-type and mutagenized MscS in POPC [13, 14], as well as of a closed state, using a coarse-grained representation of the system (**Table 1**). The experimental structures (blue cartoons) are overlaid with calculated 3D density distributions mapping the morphology of the alkyl chain double layer in each of the MD trajectories (gold volumes), up to 10 Å from the protein surface. Maps are represented as in **Figure 4**. Data represent averages over 20 μs of simulation.
